# Visual Analytics for Deep Embeddings of Large Scale Molecular Dynamics Simulations

**DOI:** 10.1101/830844

**Authors:** Junghoon Chae, Debsindhu Bhowmik, Heng Ma, Arvind Ramanathan, Chad Steed

## Abstract

Molecular Dynamics (MD) simulation have been emerging as an excellent candidate for understanding complex atomic and molecular scale mechanism of bio-molecules that control essential bio-physical phenomenon in a living organism. But this MD technique produces large-size and long-timescale data that are inherently high-dimensional and occupies many terabytes of data. Processing this immense amount of data in a meaningful way is becoming increasingly difficult. Therefore, specific dimensionality reduction algorithm using deep learning technique has been employed here to embed the high-dimensional data in a lower-dimension latent space that still preserves the inherent molecular characteristics i.e. retains biologically meaningful information. Subsequently, the results of the embedding models are visualized for model evaluation and analysis of the extracted underlying features. However, most of the existing visualizations for embeddings have limitations in evaluating the embedding models and understanding the complex simulation data. We propose an interactive visual analytics system for embeddings of MD simulations to not only evaluate and explain an embedding model but also analyze various characteristics of the simulations. Our system enables exploration and discovery of meaningful and semantic embedding results and supports the understanding and evaluation of results by the quantitatively described features of the MD simulations (even without specific labels).

## I. Introduction

Molecular dynamics (MD) simulations provide meaningful insights into the atomistic details of complex biological processes such as protein folding. Under the hood MD essentially solve Newton’s laws of motions for large groups of atoms (e.g., a protein or other complex biological structure) and generate vast quantities of visually rich information that need to be analyzed quantitatively for further detailed interpretation. Therefore, data generated from MD simulations tend to be high-dimensional and with millions of data points [1]–[6]. For example, a typical protein folding trajectory can possess tens of thousands atoms (including solvent) and generate millions of conformations.

From a computational perspective, one of the major challenges is the efficient extraction of experimentally relevant features from these high-dimensional datasets. These features often tend to be low-dimensional. Given the inherent high dimensionality of MD simulation datasets, it is often necessary to use dimensionality reduction methods to relate these datasets with experimentally relevant features. Dimensionality reduction techniques result in latent embeddings of the MD data such that they can be compared with experimental data; however, this is still challenging since interaction with such high-dimensional datasets can pose inherent cognitive and visual challenges to elucidate biologically meaningful features.

One of the popular approaches to evaluate the embedding models and analyze the underlying features is to visualize them. The interactive visualization of these embeddings allows one to not only verify dimensional reduction methods (i.e., how models accurately capture certain similarity across groups of simulation frames), but also potentially interpret bio-molecular mechanisms that lead to specific observations across MD simulations [7]–[9]. However, most existing visualizations for embeddings have some limitations in evaluating the embedding models and understanding the complex MD simulations. Such visualizations are non-flexible and non-interactive, and provide limited functions. Embedding projector [10] demonstrated an interactive visualization system for embeddings and analysis of high-dimensional data to mitigate the limitations. Although the system is designed for general purposes, the system still has some limitations related to user interactions and analysis for data with no labels.

In this paper, we propose a new visual analytics system for embeddings of MD simulations to not only evaluate and explain the embedding model but also observe various characteristics of protein conformational changes. Our system allows scientists to explore and discover meaningful geometry of conformational state clusters. Existing visual representations for embeddings usually attach metadata (labels) to the data points in the latent space to understand the models. Although our MD simulation data do not have specific labels, it can be quantified by specific measurements, such as the total number of contacts, the fraction of native contacts, and root mean squared deviations (RMSD) to the native state. Our system supports the understanding and evaluation of the clustering results by coordinating the quantitatively described features of the MD simulations. Also, we provide the improved ability of user interaction for effective visual analysis of embeddings. Finally, the system allows a in-depth analysis of the MD simulations by rendering and visualizing the molecules.

## II. Related Work

Several novel approaches for visualizing outputs from machine learning and deep learning have been proposed [11]. However, we discuss previous work related specifically to the visualization of latent dimensions (i.e., embeddings) from dimensionality reduction techniques [10], [12]–[15]. Embedding Projector [10] provides an interactive 2/3D projection for embeddings of high-dimensional data using PCA and t-SNE [16]. ACTIVIS [12] also provides a projection view of instances; however, it is more focused on visualizing how neurons (within deep learning algorithms) are activated by instances or instance subsets to understand how a model makes decisions. EmbeddingVis [13] provides a visual analytics system for exploratory and comparative analysis of graph embeddings models. Some studies [14], [15] have focused on visualizations for exploring and analyzing word embeddings. While these previous studies can be generally used for visualization, they do not necessarily address scientific domain specific (i.e., in this case, molecular dynamics simulations) datasets and how biophysically relevant insights can be drawn by just visualizing these datasets.

In this paper, we enable new user interactions with visualizations for embeddings in 3D space and demonstrate how our system handles embeddings of long time-scale molecular simulation datasets with quantitative descriptions rather than qualitative labels. Finally, to the best of our knowledge, this is the first interactive visualization for deep embeddings of MD simulations.

## III. Embedding of Molecular Dynamics Simulations

### A. Background

Proteins perform their physiological functions by structural transitions between various native conformational states through folding/unfolding or association/dissociation with ligands or other bio-molecules. With the extensive application of GPU, MD simulation can now sample trajectories (i.e., structural protein transitions) up to experimentally relevant millisecond time scales, capturing the dynamics in atomistic detail. To elucidate the mechanism and pathway of protein structural transitions, therefore, it is important to observe, identify, and characterize local conformational states from the MD trajectories. However, the long-timescale MD simulations consequently generate a massive amount of high-dimensional data in size of terabytes. Current analysis methods are often challenged in handling these kinds of data. Therefore, recent studies have been seeking to analyze MD simulations using high-performance computing to reduce the data dimensionality and capture the most important and effective dimensions from the simulation trajectories for such biological processes [1], [6], [17]. Thus, many dimensionality reduction techniques have been developed and employed to embed the 3 × *N* (*N* is the number of atoms in the molecule) dimensional conformational states in a low-dimensional latent space (2D or 3D) [18], [19]. The result embeddings are commonly projected into a 2D or 3D plot. But it often takes elaborating efforts to extract an individual point or a subset of points for further analysis. In this paper, we introduce a new visual analytics system for the embeddings, which enables users to interact with the MD conformers in the latent space.

### B. Dimensionality Reduction

The dimensionality reduction process can be conducted using an autoencoder, a self-supervised deep learning model, which can be trained to reconstruct a dateset after encoding it into a reduced latent space [20]. Various autoencoders have been developed and successfully constructed lowdimensional underlying representations of the 3D protein conformations [7]–[9]. In this study, an application of a convolutional variational autoencoder (CVAE) was adapted to automatically reduce the high dimensional conformations from MD simulations into scattering points in 3D latent space where those points are also grouped according to shared structural and energetic characteristics [7], [8], [21]–[24]. Instead of a fully-connected autoencoder learns an arbitrary function, we use a variational autoencoder (VAE) that encodes the inputs as a normal probability distribution function that in latent space. As a result, the data points from VAE are more evenly distributed in the latent space, which leads to a better visualization outcome. Furthermore, convolutional layers are added to the VAE architecture to utilize their sliding filter maps that can take account neighboring information into each point in the contact maps. Because conformational information in protein appears as local patterns in contact maps, convolutional layers are better suited to recognize these patterns comparing to feedforward networks. With the trained model, the conformational information from MD simulations are embedded into a latent space spanned by the three dimensions (*z*_1_ – *z*_3_). Each embedding corresponds to a conformational state, and thus the folding reaction coordinates of protein folding/unfolding process [25].

### C. Metadata

To gain better visualization of the latent data, the points in the latent space are painted with different colors according to features of the corresponding conformations in real space. While MD trajectories contain only the atom positions at the different time frame and are lack of inherent labels to define such features, each data point can be labeled with the results from quantitative analysis, which can distinguish the individual protein conformation. The commonly used methods are the total number of contacts, the fraction of native contacts, and RMSD to the native state. Native contacts are based on a cutoff distance of 8 A between C_α_ atoms. To calculate the fraction of native contacts we use a definition from the work of Savol and Chennubhotla [26]. The RMSD of each conformation are calculated against the fully folded native state and at least 75% of conformations remain within an RMSD cut-off of 1.1 Å of the native state.

**Fig. 1.**
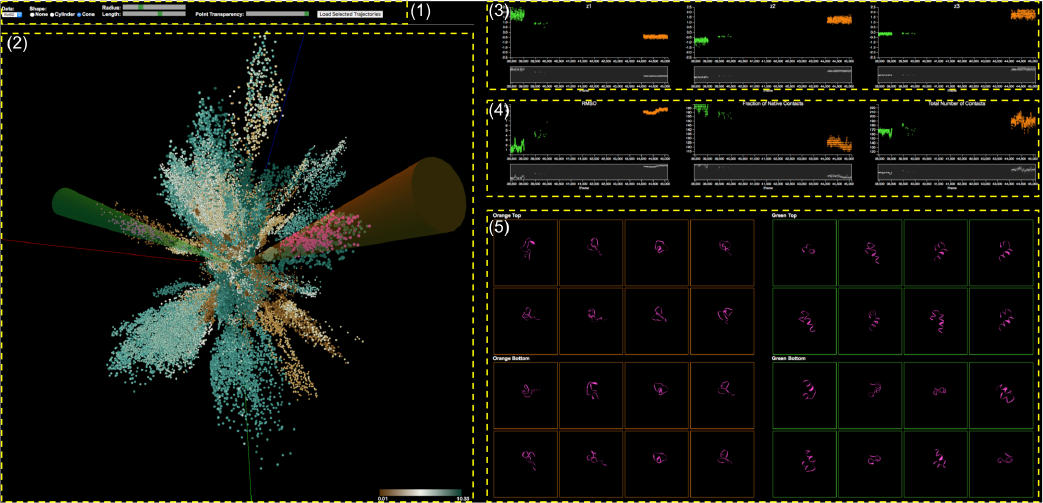
Analysis of MD simulation embeddings of a visualization system with the control panel (1), the 3D Embedding View (2), the Individual Dimensional Component View (3), the Metadata View (4), and the Molecular View (5).

## IV. System Design

The overall goal of analyzing embeddings of MD simulations is to understand biochemical processes such as conformational changes and then get insights into such bio-molecular mechanisms. To achieve the goal, multiple requirements should be achieved. We identified three design requirements:

- **R1. Evaluation**: Scientists need to evaluate, explore, and compare the clusters of conformational states by the embedding models.
- **R2. Interpretation**: Since the conformational states have no specific labels, we need to provide an easy way to distinguish and interpret the states.
- **R3. Examination**: Analysts need to view the actual conformational states (molecules) of interest for verification and further examination.

With respect to these data specifics and the requirements, our visualization system provides suitable visual representations and highly interactive features in different views for analyzing embedding of MD simulations. In Figure 1, our visualization system consists of multiple components: the control panel (1), the 3D Embedding View (2), the Individual Dimensional Component View (3), the Metadata View (4), and the Molecular View (5). All views are connected with brushing and linking. These multiple coordinated views enable scientists to explore different aspects of the data in different views through different representations so that they find causal relationships easier and uncover unforeseen connections. The following sections describe each view and user interactions with the views.

### A. 3D Embedding View

After dimensionality reduction, we are given a set of data points in the 3D latent space. Our 3D embedding view displays the 3D data points as circle dots as shown in Figure 2. The position of each dot represents its *z*_1_, *z*_2_, and *z*_3_ coordinate.

**Fig. 2.**
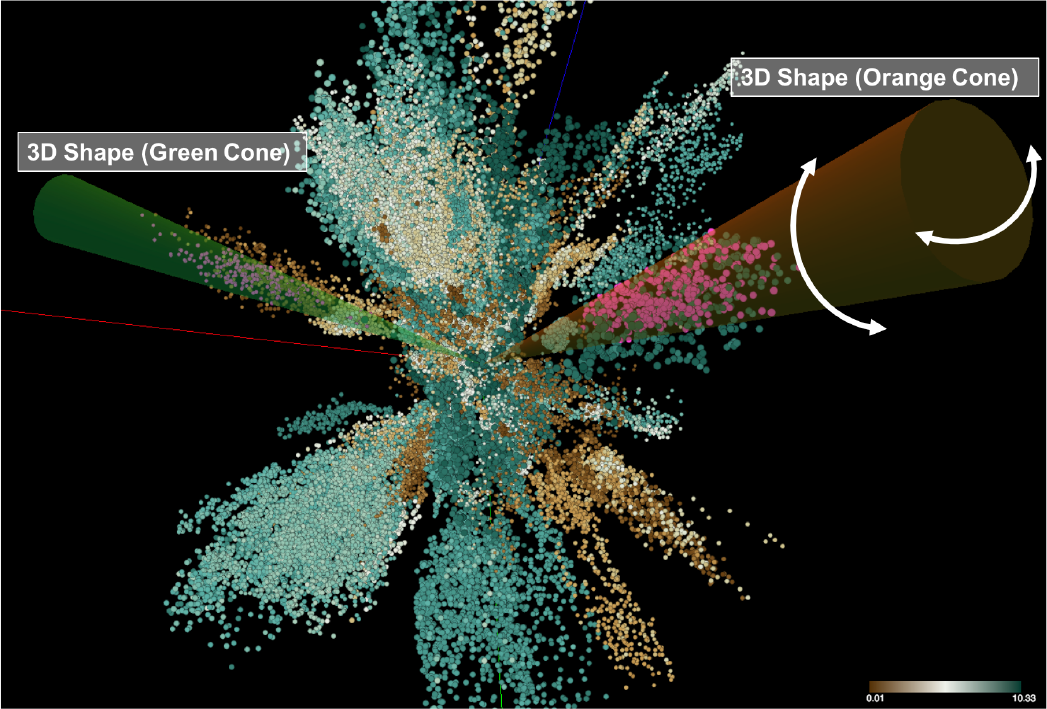
3D Embedding View: 3D scatter plot for embeddings of MD simulation and interactive cluster selection by shape templates.

The color of each dot encodes the value of the selected metadata (RMSD selected). Users are able to choose one of the metadata sets. Data points with high values are colored by dark green and low values are dark brown. Users easily explore the data points embedded into the 3D space to see which of them are close together and what are their values. We can see that some data points are closely placed to each other in an elongated shape which is a cluster of conformational states, where we can clearly see that each cluster has its own direction. As mentioned earlier, the value of metadata represents a specific characteristic of the conformational state. Therefore, users can evaluate their embedding model; how the data points are separated into distinct clusters; how the colors of points in a cluster are similar to each other (R1, R2).

Scientists often want to select a specific cluster to investigate it in detail. Also, they want to compare two different clusters to identify different characteristics (i.e., folded/unfolded). Our system allows them to select clusters using two shape templates that have two different types available: cylinder and cone, where the shapes serve as filters. They select one of the types. If they select *Cone*, two 3D cone shapes with different colors (Green and Orange) appear in the 3D embedding view (See Figure 2). If they select None, the shapes disappear. Once the selected 3D shapes appear, they can rotate each shape in any direction. They select a shape by clicking on one of the shapes and rotate it by mouse moving like a joystick, where the origin of the shape is anchored to the center of the view. Clicking on the view outside of the shapes enables rotating the camera position (viewpoint) of the 3D view. In addition to rotating the shapes, they are able to change their length and radius. The selected data points that are inside the shape are colored pink (See Figure 2). This selection task is very important for interpreting and understanding the evolving process of conformational states since each cluster represents a specific conformational state. This interactive visualization enables the continuous representation of the selected clusters of interest and rapid, incremental feedback for the users which allows them to complete the selection task in less time because they can see and evaluate the results of an action before finishing the action [27].

**Fig. 3.**
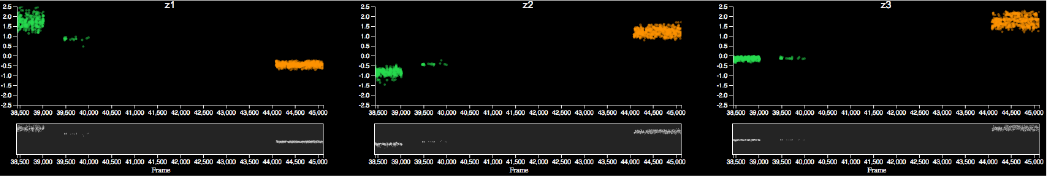
Individual Dimensional Component View: The three components of each selected data point are displayed on the three scatter plots separately: *z*_1_ (left), *z*_2_ (center), *z*_3_ (right). Each plot consists of two plots: focus (top) and context (bottom) plots. The colors of dots in the focus plot represent the corresponding 3D shapes.

The rationale behind such a design is that the 3D scatter plot is the better choice than a 2D scatter plot. We also conduct experiments with the embedding model to reduce the high dimensional data to a 2D latent space. The results of experiment show that the clusters are too severe overlap to visually distinguish. Also, it is hard for users to interactively select a single cluster. So, even though the 3D plot has many limitations, in this study it provides better performance than 2D. Also, as users can easily rotate the viewpoint of the 3D scene, the occlusion issue is mitigated.

### B. Individual Dimensional Component View

Interpreting each dimensional components is very important for understanding the embedding models. For example, the model developers may want to know which components are strongly correlated with a specific cluster and which components have high variation. Once scientists select two sets (clusters) of the data points in the 3D embedding view, the three components (*z*_1_, *z*_2_, *z*_3_) of the each selected data point are separately displayed in the three different plots, visualizing the distributions of each dimensional component (R1). The three plots are lined up horizontally in order *z*_1_, *z*_2_, and *z*_3_ (Figure 3). The x-axes are the frame number (the temporal sequence of conformational states). As we can see, however, a large number of points causes a clutter issue. To resolve the issue, we employ a focus+context interface through brushing [28]–[30]. Each plot consists of two sub plots: focus and context plots. The context plot (bottom) represent the overall distribution of the entire selected data points. The focus plot (top) shows only a subset, where the subset is selected by brushing over a specific region of the context plot. Also, the plots are linked together, so if users brush a specific area on one of the context plots, the same areas of other plots are brushed. In Figure 3, the plots show the individual dimensional components of the selected clusters shown in Figure 2. The green and orange dots of the plots indicate the selected clusters by the green and the orange cone shapes respectively. The cluster (green dots) has a higher variation on the *z*_1_, while another one (orange dots) has a higher variation on the *z*_3_ (see Figure 3).

**Fig. 4.**
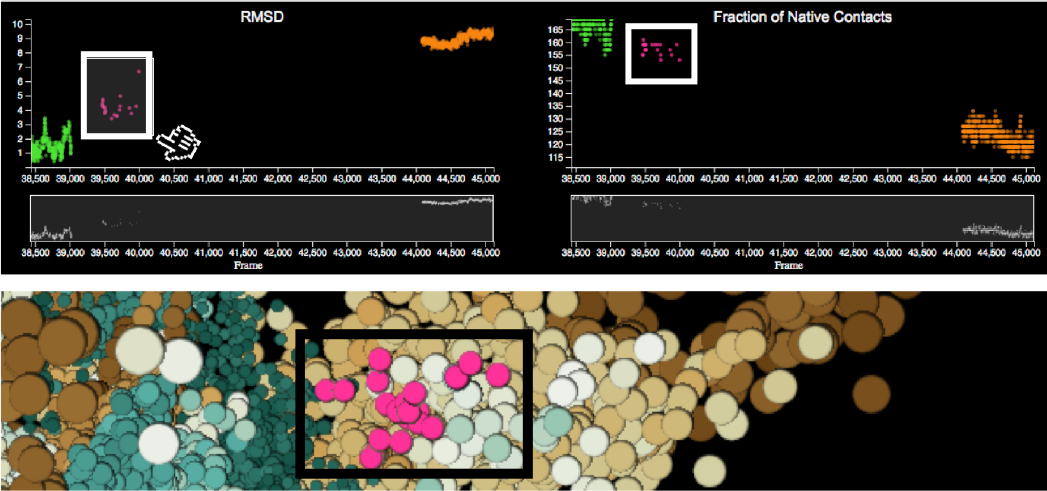
Metadata View: Each plot shows the different metadata of the selected data points (top). The plots and the 3D embedding view (bottom) are coordinated.

### C. Metadata View

Although the conformational state data is highly complex, each state can be quantitatively described by corresponding metadata described earlier. Our metadata view provides three plots for each metadata item to provide a straightforward way to interpret and classify the states (R2). In Figure 4, we show two plots for RMSD (top-left) and the fraction of native contacts (top-right) out of three plots as an example. The plots are also designed in the same manner as other plots. Also, scientists are able to select points of interest through brushing on the focus plot. The selected points are colored pink in all the plots. The 3D embedding view is also coordinated through the synchronous highlighting of corresponding elements, where the elements are colored pink as well (See Figure 4 (bottom)). These fully linked views enable users to explore different aspects of the data in different views and to find causal relationships easier.

### D. Molecular View

Our molecular view allows scientists to view and interact with actual 3D MD simulations of the selected data points (R3). Since the size of the selection is usually very huge (1K – 3K), we sample the data points based on the point’s magnitude in the direction of each cluster. Each cluster has a certain direction (vector from the origin to the center of the shape’s bottom). For each selected point, we calculate its scalar projection of the point’s vector onto the cluster’s vector. Then, we choose two sets of the points: eight points with the highest values and eight points with lowest values to view balanced samples.

Snapshots of a specific viewpoint of the selected MD simulations are displayed in the small square views initially in Figure 5. Figure 5 shows some sample views of all views (5) in Figure 1. Once they click on one of the views, it is converted into a 3D viewer for MD simulations (see first and third ones in Figure 5). The views enable users to interact with the actual 3D structure of the corresponding MD simulation. Then, they can rotate and zoom the 3D structure using the viewer. Also, hovering the mouse cursor on it shows detailed information of the MD simulation. In Figure 5, the first and second simulations in the green boxes show two samples respectively as examples out of the simulations selected by the green cone in the 3D view. The third and last ones are selected by the orange cone. The first and third simulations represent the data points with high scalar projection value; the second and last ones do the data points with low value. The simulations in the green boxes correspond to the green dots with low RMSD and high fraction values in the two metadata plots in Figure 4. Through the 3D structures we can verify that the protein chains are folded (helix structure) (R3). The ones in the orange boxes correspond to the orange dots with high RMSD and low fraction values. In Figure 5, we can see that the protein structures are unfolded.

**Fig. 5.**
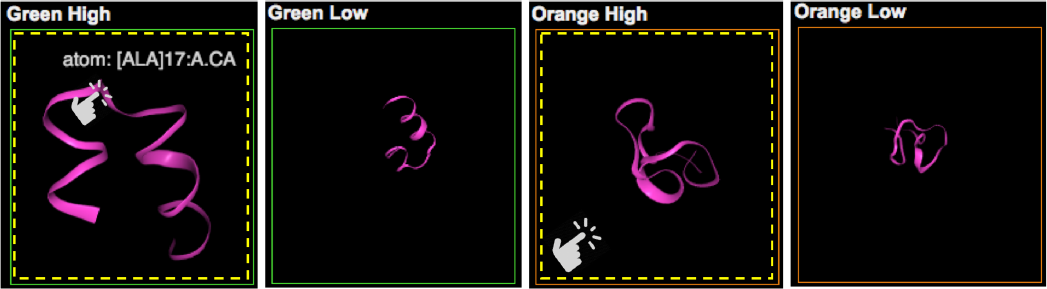
Molecular View: Each small view represents an actual MD simulation. Clicking on one of them, it is converted a 3D viewer for MD simulations, which enables interactive analysis of its 3D structure.

The rationale of this visualization design is that the domain experts need to see the actual 3D structure from the different viewpoints of the selected data points for verification and further examination. Without our system, for this work, they have to use another tool to load and view the 3D structure for each data point. This task is burdensome and ineffective.

## V. Case Study

This study is still on an initial stage. We work with three domain experts on a pilot case study. Here, we emphasize how our visualization system enables exploration and identification of meaningful embedding results and how the system can help the experts evaluate their dimensionality reduction model.

The dataset consists of 28 separate MD trajectories of the Fs-peptide, a widely studied model system for protein folding. To demonstrate the system, MD simulation results of Fs-peptide are prepared through dimensionality reduction and computation of metadata. For protein folding, resulting in an aggregate sampling of 14 *μ*s, the dataset is consisting of 280,000 conformations (data points). We processed each conformations using the MDAnalysis library [31], [32] to extract contact maps between every pair of *C_α_* atoms; we consider an atom to be in contact to another atom if it is separated by less than an 8 Å. Contact map provides a reduced representation of protein structure from its full 3D atomic coordinates. Note that contact map representation is invariant to rotation and translation (which is typically an artifact of MD simulations). The contact maps are successively fed into our dimensionality reduction architecture.

We apply the convolutional variational autoencoder to reduce the high dimensional conformations from MD simulations into 3D latent space. In Figure 2, we can see there are many clusters and particularly the data points of each cluster have similar RMSD values (the higher the RMSD value, the more unfolded the state. The lower the RMSD, the more folded the state). We can observe that different states are separated into distinct clusters. Also, it is notable that the folded states (brown) are similarly clustered together.

**Fig. 6.**
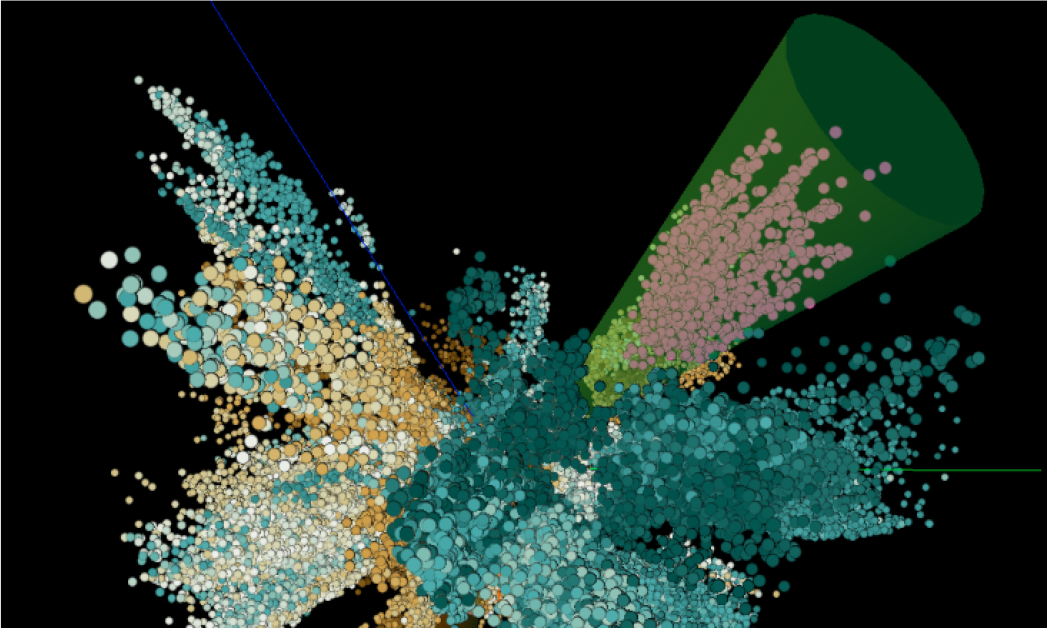
Cluster Selection: A set of points are selected by a 3D cone shape to investigate the embedding model and the cluster in detail.

**Fig. 7.**
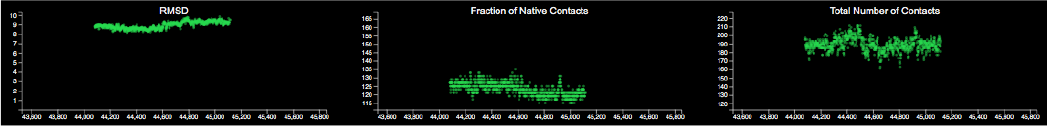
Cluster Analysis: Each plot shows three different metadata values of the selected cluster in Figure 6.

We evaluate the model, how it classifies the protein conformation based on the corresponding metadata, inherent characteristics of conformations. We find out two metadata values tend to have an inverse correlation through the two plots: RMSD (top-left) and the fraction of native contacts (top-right) in Figure 4. We also select another cluster using the green cone shape (see Figure 6). In Figure 7, the three metadata plots show each corresponding metadata. The cluster has very similar RMSD values (left) while there is a larger variation on the fraction of native contacts (center) and the total number of contacts (right). Similar results are shown even when we select other clusters. Therefore, we can assess the current model provides better results in terms of the RMSD than other metadata.

## VI. Conclusion

We proposed a new visual analytics system that analyzes the embedding of MD simulations by a dimensionality reduction framework. The system also helps scientists interpret the complex bio-molecular mechanisms using interactive and coordinated visualizations. Selecting the clusters in the 3D Embedding view using a mouse is still tricky. As future work, we will improve the user interaction to improve the operation. Also, we will allow the system to compare multiple embedded models to find an optimized model.

## Acknowledgment

This material is based upon work supported by the U.S. Department of Energy, Office of Science, Office of Advanced Scientific Computing Research, under contract number DE-AC05-00OR22725. This research is sponsored in part by the Laboratory Directed Research and Development Program of Oak Ridge National Laboratory, managed by UT-Battelle, LLC, for the U. S. Department of Energy.

This research used resources of the Oak Ridge Leadership Computing Facility at the Oak Ridge National Laboratory, which is supported by the Office of Science of the U.S. Department of Energy under Contract No. DE-AC05-00OR22725.

